# Body-segment coordination as a new predictor of freezing of gait in people with Parkinson’s disease

**DOI:** 10.64898/2026.02.02.702843

**Authors:** Marco Romanato, Antonio Carlos Costa, Anthony Jannou, Anqi Zhou, Saoussen Cherif, Elodie Hainque, David Maltête, Stéphane Derrey, Mathieu Yeche, Carine Karachi, Claire Wyart, Brian Lau, Marie-Laure Welter

**Affiliations:** Sorbonne université, Institut du Cerveau-Paris Brain Institute-ICM, Inserm U1127, CNRS UMR7225, APHP, F-75013 Paris, France; Neurology Department, Hôpital Pitié-Salpêtrière, AP-HP, F-75013 Paris, France; Neurology Department, CHU Rouen, Rouen University, F-76000 Rouen, France; Neurosurgery Department, CHU Rouen, Rouen University, F-76000 Rouen, France; Neurosurgery Department, Hôpital Pitié-Salpêtrière, AP-HP, F-75013 Paris, France; Neurophysiology Department, CHU Rouen, Rouen University, F-76000 Rouen, France

**Keywords:** Parkinson’s disease, freezing of gait, limb coordination, subthalamic nucleus, deep-brain stimulation

## Abstract

Freezing of gait (FOG) in Parkinson’s disease (PD) involves impaired integration of posture and locomotion with altered body-segment coordination. We measured coordination using gait kinematics during walking in 16 PD patients with FOG, preoperatively with and without dopaminergic medication (OFF/ON-DOPA), and postoperatively with and without subthalamic deep brain stimulation (OFF/ON-DBS). Body-segment coordination was modeled from acceleration-based intersegmental correlations across trunk, pelvis, and limbs. We tested whether coordination metrics predict individual postoperative FOG severity using LASSO regression with nested cross-validation, including preoperative demographics, clinical scores, gait and coordination metrics. Preoperatively, DOPA decreased trunk-pelvis-upper-limb coordination but increased crossed upper-lower-limbs coupling; while STN-DBS selectively increased inter-upper-limb coordination, with DOPA- and STN-DBS-induced changes being correlated. Whole-body coordination predicted individual postoperative FOG severity, with the most important couplings being the trunk-pelvis and pelvis-lower-limb. Body-segment coordination captures clinically relevant gait metrics in PD, highlighting coordination as a potential biomarker for patient stratification and treatment response.

## Introduction

Body-segment coordination is crucial for natural displacement in the environment, enabling the integration of posture and movement to adapt to anticipated and unexpected changes, such as obstacles, directional adjustments, or social interactions. Across vertebrates, body-segment coordination is closely linked to the mode of displacement. In anamniote species, swimming relies on head-to-tail coordination^1–3^. In mammals, locomotion is limb-based rather than undulatory, relying on right-left alternation and rostro-caudal sequencing of limb movements^4^. In humans, bipedal upright posture adds an additional level of complexity, requiring continuous coordination between lower-limb (LL) movement and postural control to ensure smooth transitions between static and dynamic walking phases^5^.

Neural structures involved in limb pattern coordination are partially conserved across species and include spinal central pattern generators (CPGs), the mesencephalic locomotor region (MLR), sensory feedback, and intersegmental coupling^3,6^, and the cortico-basal ganglia networks, including the supplementary motor area (SMA), which connects with the contralateral motor cortex^7^. A tight functional coupling between the MLR and reticulospinal-CPGs pathways plays a critical role in gait initiation, modulation of locomotor frequency, postural adjustments, and LL coordination^8,9^. In humans, an additional level of coordination exists between upper (UL) and LL during walking, with UL movements leading LL patterns^10–12^. This organization supports the concept of a dual-oscillator system for arm-leg coordination mediated by long propriospinal neurons^13^.

In Parkinson’s disease (PD), the adaptive integration of posture and locomotion is progressively impaired. One of the earliest motor hallmarks of PD is slowed walking with reduced arm swing, reflecting nigrostriatal dopaminergic degeneration and basal ganglia dysfunction associated with less efficient neural coding^14–16^, and a progressive disruption of intersegmental functional connectivity within the spinal cord^17^. As the disease progresses, postural instability, falls, and episodes of freezing of gait (FOG) emerge^18,19^. FOG is characterized by sudden, episodic arrests of forward walking, frequently triggered by gait initiation, turning, or dual-task situations, and by a functional disconnection between preparatory programming and motor execution^19–22^. Altered coordination during walking has also been reported in PD patients with FOG^23–25^. Discoordination of the LL, with disrupted reciprocity, timing and muscle activation pattern, has been described and worsens under complex walking conditions^26–28^. Coordination deficits are not restricted to the LL with motor blocks of the UL during rapid alternating movements^29^, and discoordination of all four limbs has also been reported during tasks such as crawling, swimming or dancing^30,31^. In this context, increased UL-LL coupling could reflect a compensatory strategy aimed at stabilizing gait by reducing cortical drive when automatic interlimb coordination is impaired^12,32^, highlighting the possible role of body-segment coordination breakdown in the pathophysiology of FOG.

Despite these advances, FOG remains therapeutically challenging. Its response to dopaminergic medication is often incomplete or inconsistent^22^, and deep brain stimulation (DBS) outcomes are highly variable^33,34^. DBS of the MLR has shown limited efficacy^35–37^, and optimal STN-DBS can fail to relieve, or even worsen, FOG and LL discoordination^38,39^.

Here, we aimed to assess the effects of dopaminergic medication and STN-DBS on body-segment coordination in PD patients with FOG, and to relate coordination metrics to FOG severity to identify predictors of individual postoperative outcomes. To this end, we derived a holistic representation of whole-body coordination from kinematic recordings obtained during walking in PD patients and examined its relationship with FOG severity and treatment-induced changes.

## Results

Complete preoperative (PREOP) and postoperative (POSTOP) gait kinematic datasets were obtained from 16 PD patients (see supplementary materials). In these patients, both DOPA (PREOP) and STN-DBS (POSTOP) significantly reduced the severity of parkinsonian motor disability (UPDRS III) and gait and balance disorders (GABS). Motor and gait scores were significantly lower in the ON-DOPA PREOP condition than in the ON-DBS POSTOP condition (**Figure S2**). FOG severity was also significantly reduced following STN-DBS, with a mean FOG-Q reduction of 47%. However, individual responses were highly variable, ranging from a 9% worsening to a 96% improvement in FOG-Q score (**Figure S2**).

### Effects of dopaminergic medication and STN-DBS on gait kinematic parameters

*Pace* (RPC1) significantly improved with DOPA PREOP (OFF vs ON-DOPA, p = 0.004) and with STN-DBS POSTOP (OFF vs ON-DBS, p = 0.014), with no significant differences between DOPA and STN-DBS (p=0.617, **Figure 2C**).

*Rhythm* significantly improved with DOPA PREOP (OFF-vs ON-DOPA, p < 0.001) but was not significantly modified by STN-DBS POSTOP (OFF vs ON-DBS, p = 0.0684). Nevertheless, *rhythm* did not differ significantly between the ON-DOPA PREOP and ON-DBS POSTOP conditions (p = 0.068, **Figure 2C**).

In contrast, *dynamic balance* significantly worsened with DOPA PREOP (OFF-vs ON-DOPA, p = 0.037). No significant change in *dynamic balance* was observed with STN-DBS POST-OP (OFF-vs ON-DBS, p = 0.753), but *dynamic balance* scores were significantly better ON-DBS POST-OP than in the ON-DOPA PREOP condition (p = 0.037, **Figure 2C**).

*Gait variability* and *gait asymmetry* were not significantly modified by DOPA (PREOP) or STN-DBS (POSTOP) (p > 0.05 for all comparisons, **Figure S3**).

### Effects of dopaminergic medication and STN-DBS on body-segment coordination

Across the 6 anatomical groups, 15 pairwise body-segment coordination features were derived (**Figure 1, Figure S4**). No significant changes were observed in the whole-body coordination score either PREOP between OFF- and ON-DOPA conditions (p = 0.094), POSTOP between OFF- and ON-DBS conditions (p = 0.655), or when comparing ON-DOPA PREOP and ON-DBS POSTOP conditions (p = 0.094, **Figure 3A**).

**Figure 1.**
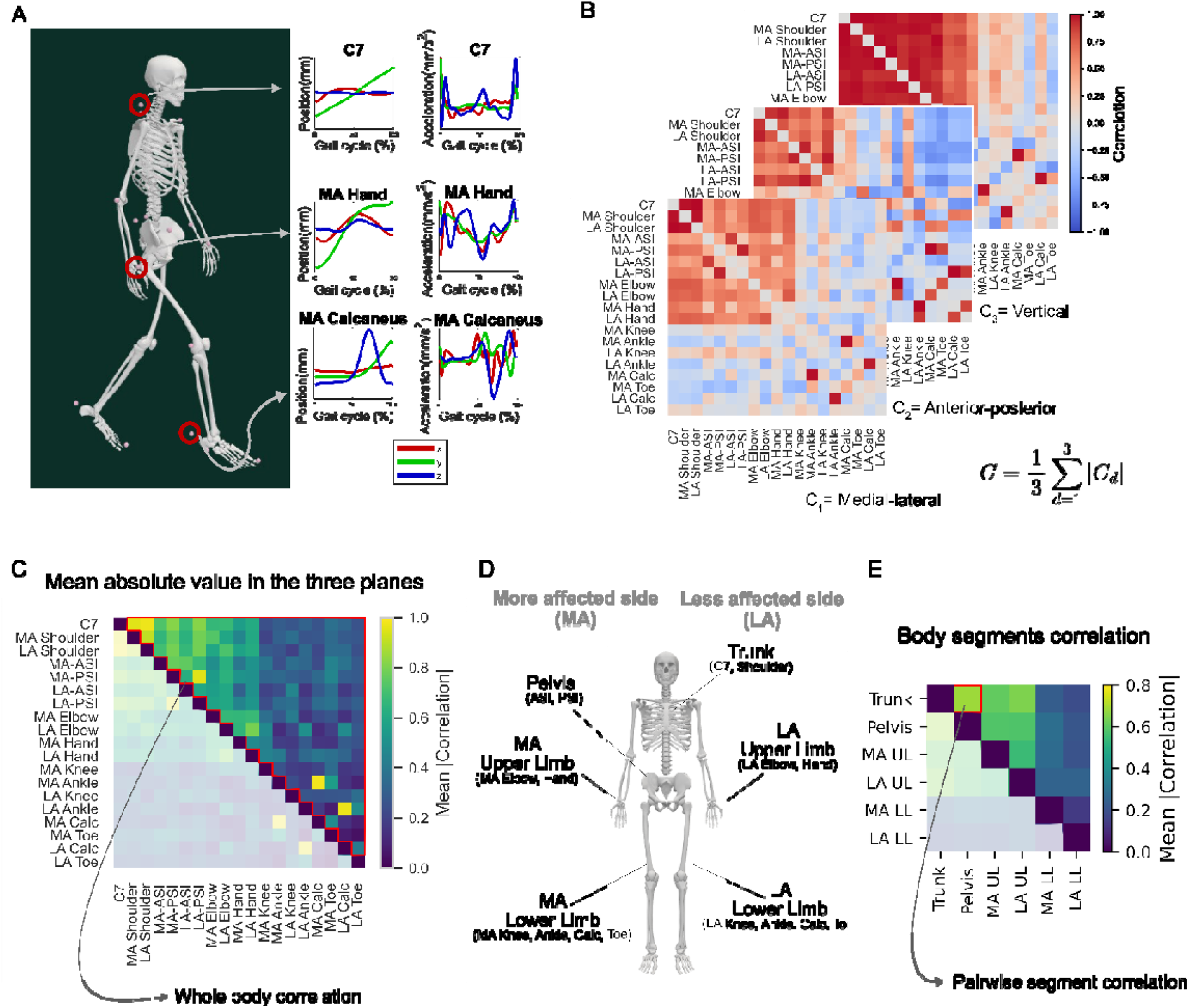
Body-segment coordination modeling. **A**. Schematic of the marker set. The mediolateral (x), anteroposterior (y), and vertical (z) position trajectories of 19 markers (*i*.*e*., the 7^th^ cervical vertebra, shoulders, elbows, wrists, anterior and posterior iliac spines, knees, ankles, calcanei, and toes) were captured, and their acceleration time series during the gait cycle were estimated (normalized by the temporal length of the gait cycle for visualization purposes). **B**. Acceleration time series were linearly correlated to produce a 19x19 correlation matrix for each axis. A final correlation matrix C was obtained by averaging the absolute values of the correlation matrix elements across the three dimensions. **C**. Correlation matrix C is obtained by averaging the absolute values across the three dimensions. The average of its upper triangular matrix is used as a score of whole-body coordination. **D**. Schematic showing how markers are aggregated to define major body segments. **E**. Final body segments correlation matrix. Aggregation of a segment pair is used as a score for the pairwise correlation of body segments.

**Figure 2.**
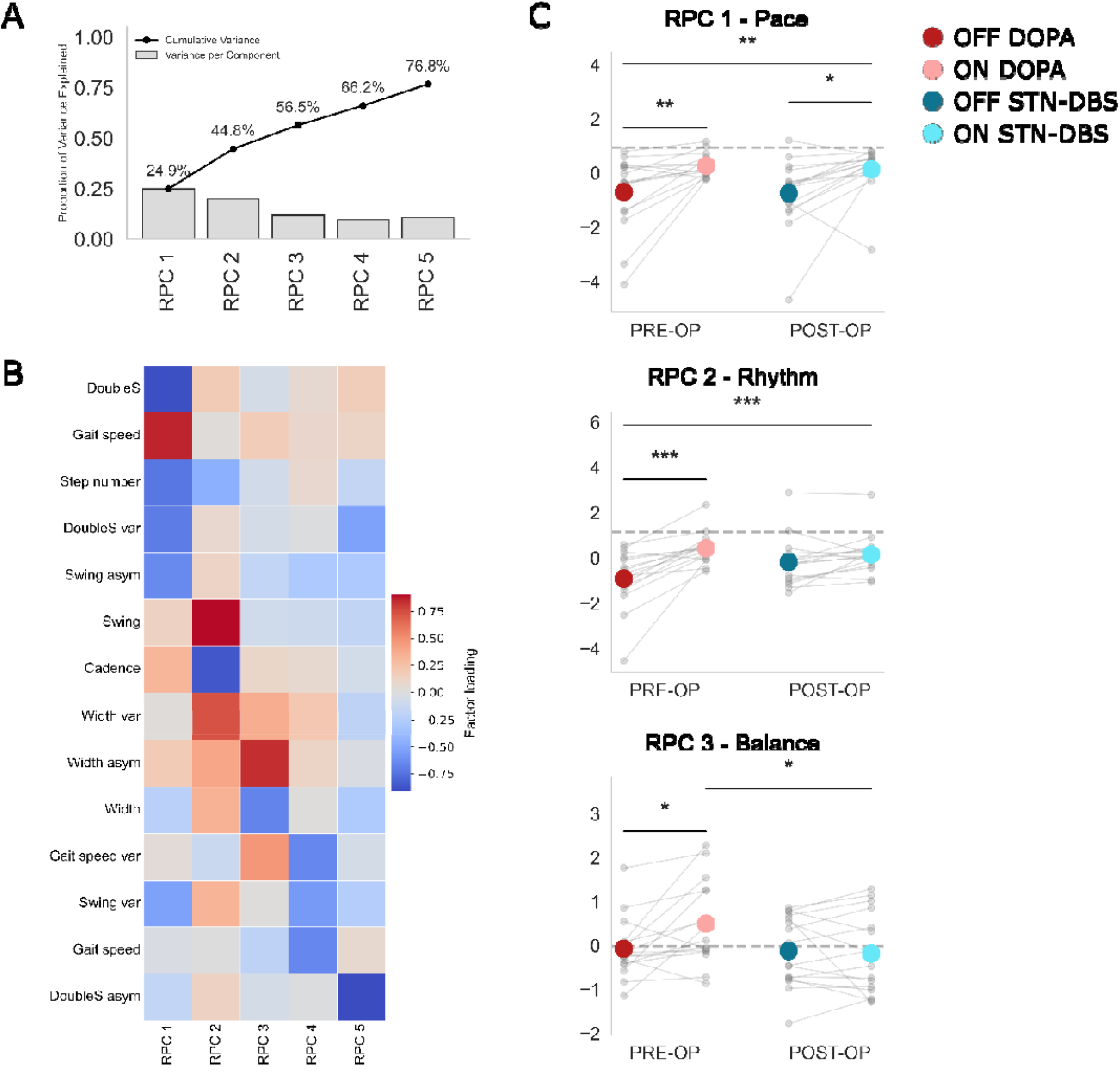
Principal components analysis on spatiotemporal parameters of walking in PD patients and effects of dopaminergic medication and STN-DBS. **A**. Variance explained by each rotated (*varimax*) component and the cumulative percentage of explained variance. **B**. Factor loadings for each component included in the RPC analysis. **C**. Graphs showing scores for *pace* (RPC1), *rhythm* (RPC2), and *balance* (RPC3) (large colored dots) obtained PREOP OFF-DOPA (red) and ON-DOPA (pink), and POSTOP OFF-STN-DBS (dark blue) and ON-STN-DBS (light blue). Each small gray dot represents one individual PD patient. The dashed gray lines represent the average of 10 healthy controls included as references. *p<0.05, **p<0.01, ***p<0.001

**Figure 3.**
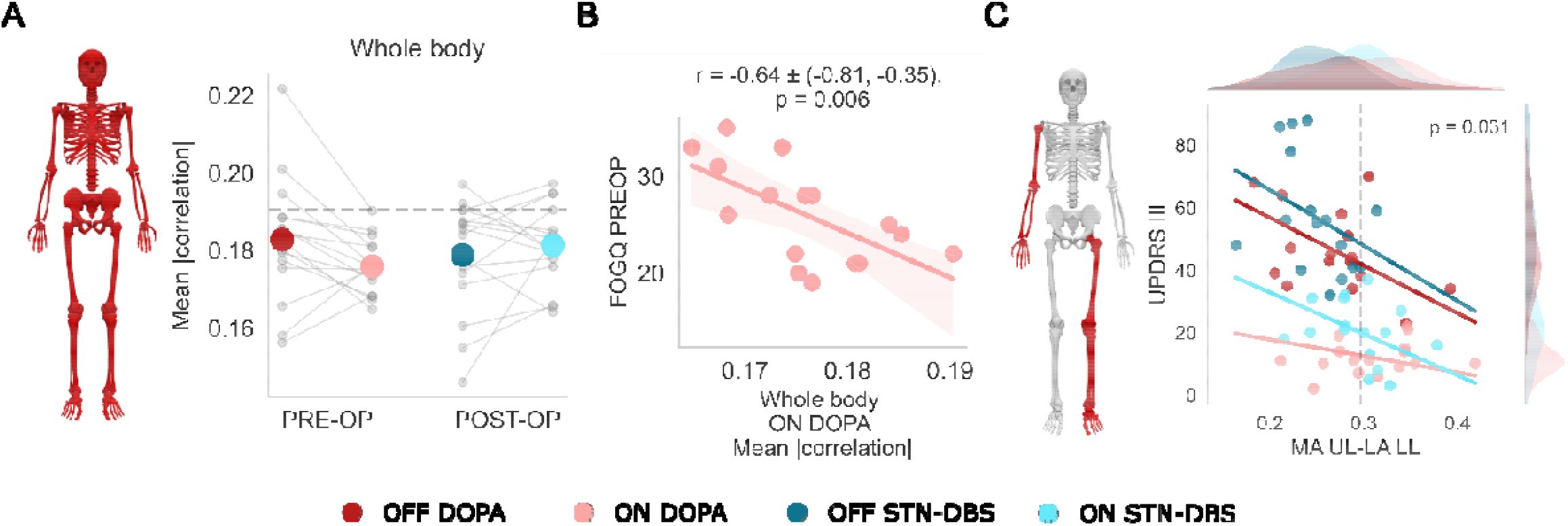
Effects of dopaminergic medication and STN-DBS on body-segment coordination in PD patients, and the relationship with FOG severity and Parkinson motor disability. **A**. The graph shows scores for whole-body coordination (large colored dots) PREOP OFF-DOPA (red), ON-DOPA (pink), POSTOP OFF-STN-DBS (dark blue), and ON-STN-DBS (light blue). Each small gray dot represents one individual PD patient. The dashed gray line represents the average of 10 HC. *p<0.05, **p<0.01, ***p<0.001 **B**. Scatterplot of the linear correlation between FOG PREOP and the whole-body coordination score ON-DOPA. Each point represents one individual. Correlation value, 95% confidence interval, and FDR-corrected p-value. **C**. Scatterplot of the linear correlation analysis between the MDS-UPDRS III scores and the “MA UL - LA LL” coordination score. Each point represents one individual, with PREOP OFF-DOPA (red), ON-DOPA (pink), POSTOP OFF-STN-DBS (dark blue), and ON-STN-DBS (light blue).

PREOP DOPA significantly decreased coordination between the trunk and MA UL (p < 0.001), the trunk and LA UL (p < 0.001), the pelvis and MA UL (p = 0.001), the pelvis and LA UL (p = 0.007), and bilateral UL (“UL MA – UL LA”, p = 0.022). Conversely, DOPA significantly increased coordination between the MA UL and LA LL (p = 0.015), between the LA UL and MA LL (p = 0.004), and between bilateral LL (“LL MA – LL LA”, p = 0.022), with no other significant changes (**Figure S4**).

POSTOP, STN-DBS significantly increased bilateral UL coordination (“UL MA – UL LA”, p = 0.029), with values significantly higher than those in the ON-DOPA PREOP condition (p = 0.008), as well as coordination between the MA UL and the LA LL (“UL MA – LL LA”, p = 0.009). This latter feature did not differ significantly from the ON-DOPA PREOP condition (p = 0.29). No other body-segment coordination features were significantly modified by STN-DBS (**Figure S4**).

When directly comparing ON-DOPA PREOP vs ON-DBS POSTOP, significant differences in coordination were observed for “Trunk – UL MA” (p < 0.001), “Trunk – UL LA” (p = 0.001), “Pelvis – UL MA” (p < 0.005), “Pelvis – UL LA” (p = 0.007), and “UL LA – LL MA” (p = 0.006), with no additional differences (**Figure S4**).

### Relationship between FOG severity and parkinsonian motor disability and body-segment coordination

We found a significant negative correlation between PREOP FOG severity and the whole-body coordination score ON-DOPA (r = -0.64, 95% CI [-0.81, -0.35], p < 0.006, **Figure 3B, Figure S5A**), with no other significant correlations.

The UPDRS III score was also significantly negatively correlated with the “UL MA – LL LA” coordination feature, decreasing 13 points per 0.1-point increase in this coordination metric (coefficient = -132.64, p = 0.031). This relationship was consistent across treatment conditions (“UL MA – LL LA” * Condition [OFF-DOPA], p = 0.778; Condition [ON-DOPA], p = 0.275; Condition [OFF-DBS], p = 0.618, compared to Condition [ON-DBS], **Figure 3C, Figure S5B**).

### Prediction of the effects of STN-DBS on FOG

Using a nested cross-validation LASSO approach with all PREOP data (demographics, clinic, gait, coordination scores) to identify predictors of POSTOP FOG severity, the model yielded R^2^ = 0.20 and RMSE = 4.77 (**Figure 4**). We found that both PREOP whole-body coordination score OFF-DOPA (3.22 ± 0.60) and GABS score ON-DOPA (2.19 ± 0.78) were retained in 100% of folds, followed by UPDRS III OFF-DOPA (-0.67 ± 0.56; 81.3% of folds) and age at inclusion (0.85 ± 0.82; 68.8% of folds). Gait domain metrics were retained in fewer than 50% of outer cross-validation folds, contributing minimally to model stability while reducing overall performance. Excluding the gait metrics from the model improved predictive accuracy (R2 = 0.66, RMSE = 3.40), and the retained predictors remained identical (**Figure S6**).

**Figure 4.**
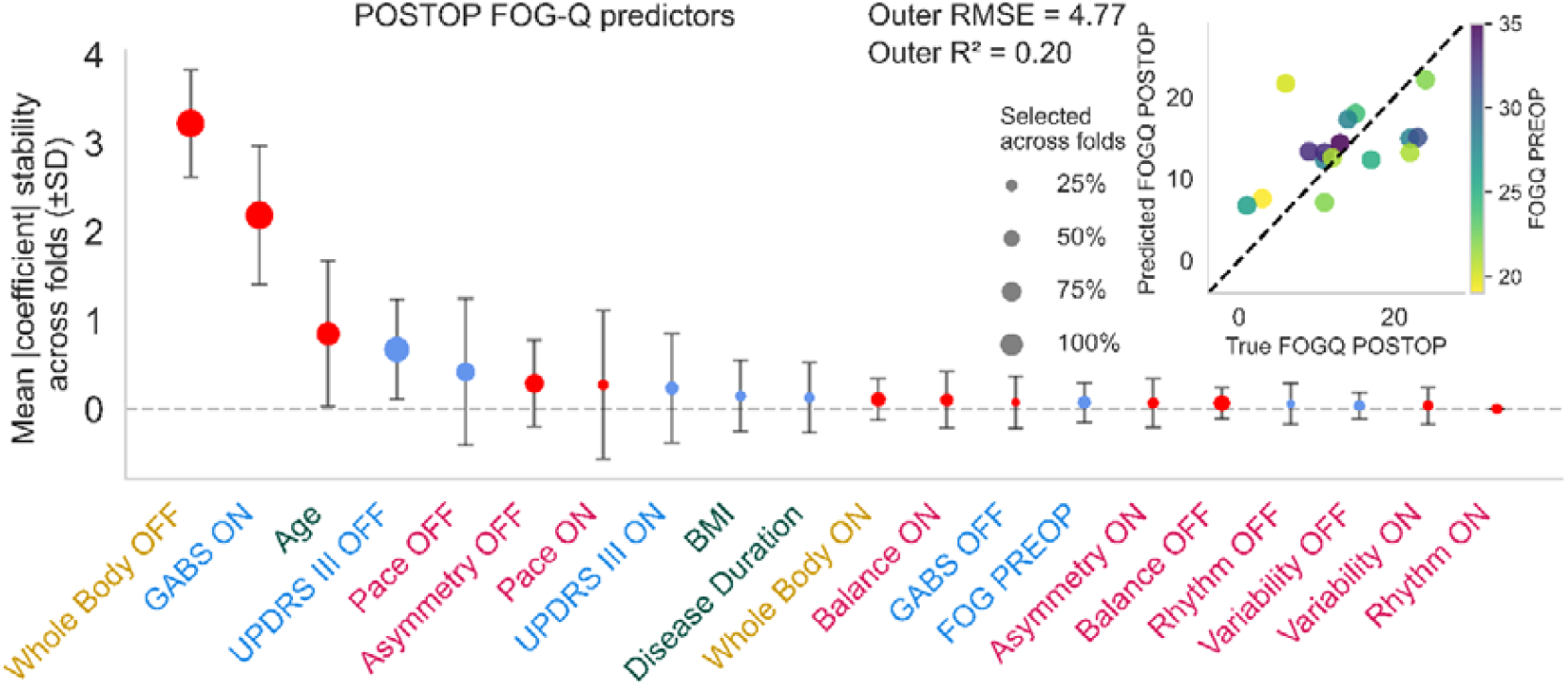
Relationships between preoperative demographic, clinical, gait domains and coordination scores, and the effects of STN-DBS on FOG severity. Coefficients stability plot of the model. Dots represent the mean coefficient across each fold, and error bars show the standard deviation, with size indicating frequency. Red dots indicate positive associations and blue dots negative associations. Out-of- fold R^2^ and RMSE are reported in the figure. X-axis labels are color-coded by feature type: coordination metrics (yellow), clinical scores (blue), demographics (green), and gait domains (magenta). The scatter plot shows true versus predicted FOG-Q POSTOP scores, color-coded by FOG-Q PREOP scores (blue: mild severity, yellow: high severity); the dashed black line represents the identity line.

Restricting the analysis to body-segment coordination metrics, four predictors were retained in 100% of outer cross-validation folds: “Trunk – Pelvis” coordination measured OFF- and ON-DOPA (OFF-DOPA coefficient = 2.92 ± 0.90; ON-DOPA coefficient = −3.22 ± 0.91), “Pelvis – LL MA” OFF-DOPA (coefficient = 4.01 ± 0.55), and “Pelvis – LL LA” ON-DOPA (coefficient = −1.77 ± 0.38), identifying the “Trunk-Pelvis-LLs” axis as a key coordination metric. This model yielded an R^2^ of 0.27 and an RMSE of 3.66 under leave-one-out cross-validation (**Figure 5A**).

**Figure 5.**
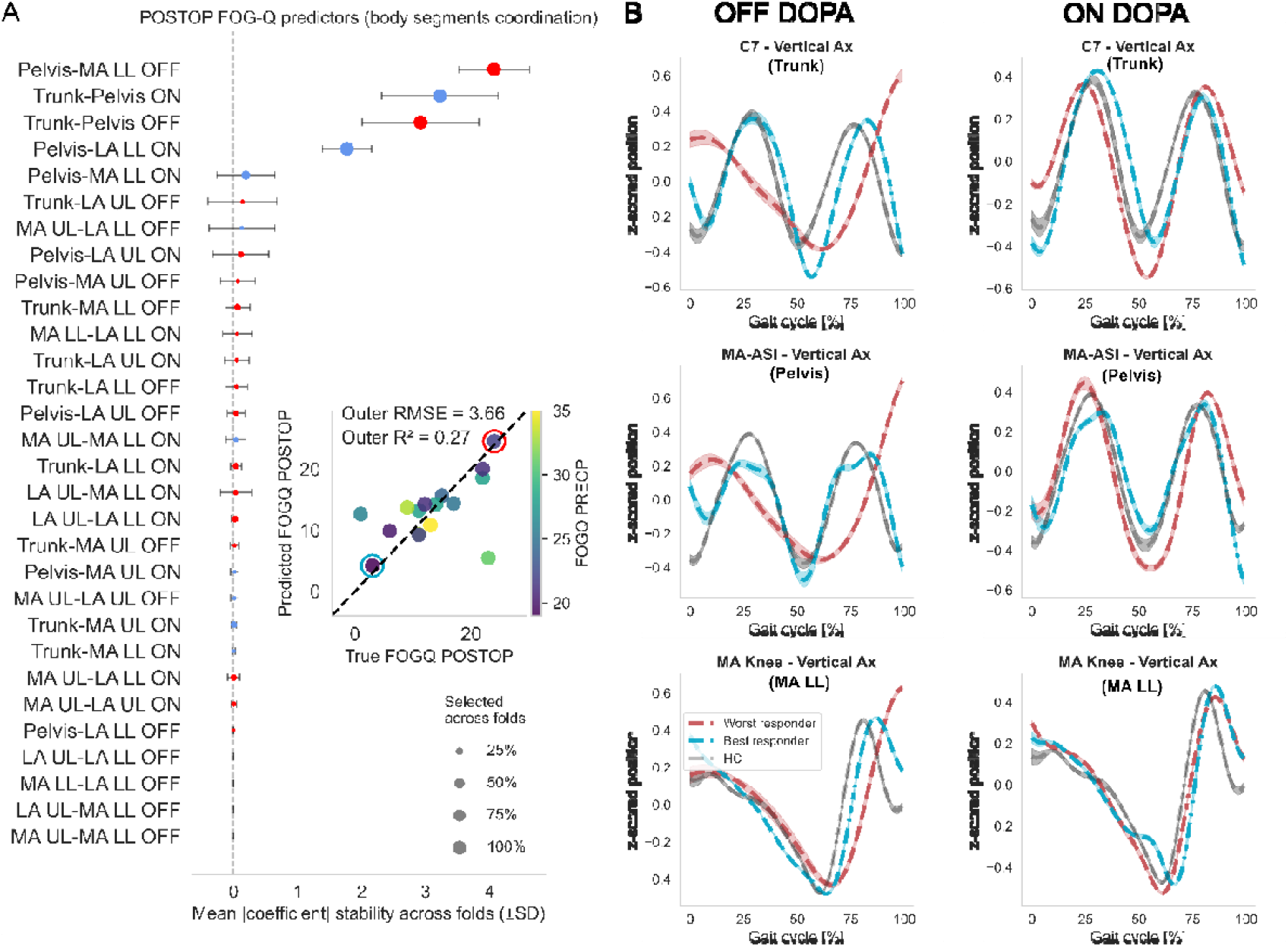
Relationship between body-segment coordination and the effects of STN-DBS on FOG severity. **A**. Coefficients stability plot of the model. Each dot represents the mean coefficient across folds between body-segment coordination scores and POSTOP FOG severity with STN-DBS. Error bars show standard deviation, and size indicates the frequency of selection across folds. Red dots indicate positive associations, and blue dots indicate negative associations. Out-of-fold R^2^ and RMSE are reported within the figure. The scatter plot shows true versus predicted FOG-Q POSTOP, color-coded by FOG-Q PREOP scores; the dashed black line represents the identity line. Best and worst responders are highlighted with blue and red circles, respectively. **B**. Position over time on the vertical plane of one representative marker per model-selected segment (‘C7’ for ‘trunk’, ‘MA-ASI’ for ‘pelvis’, ‘MA knee’ for ‘MA - LL’) for the *best* (ΔFOG-Q% = 96.2%, in blue) and *worst* (ΔFOG-Q_%_ = -9.1%, in red) FOG responders, against HS (in gray), PREOP both OFF-DOPA (left) and ON-DOPA (right). Dashed lines are the mean over each participant’s trials; shaded areas are the ± standard error. For visualization, curves were z-scored and time-normalized over 100 samples to be expressed as a percentage of the gait cycle.

Visual inspection of body-segment coordination dynamics across the gait cycle in the best responder, the worst responder, and a healthy control revealed that the best responder’s trajectory patterns closely resembled those of the healthy subject, whereas the worst responder’s patterns were markedly different. These differences were more pronounced in the OFF-DOPA condition, with partial normalization observed in the ON-DOPA condition (**Figure 5B**).

Finally, we examined the relationships between POSTOP FOG severity and PREOP DOPA effects on both whole-body and “Trunk – Pelvis – LLs” coordination, which were found to predict POSTOP FOG severity. For “Trunk – Pelvis – LLs” coordination, analyses were restricted to model-selected metrics and quantified as mean absolute values across the trunk, pelvis, and LLs segments. POSTOP FOG was predicted with high accuracy by PREOP DOPA-induced changes in both the whole-body (R^2^ = 0.48, RMSE = 4.06) and “Trunk – Pelvis – LLs” (R^2^ = 0.78, RMSE = 2.35) coordination scores (**Figure 6A**). The POSTOP effects of STN-DBS on these coordination scores were accurately predicted by PREOP DOPA effects on the same scores (whole-body: R^2^ = 0.35, RMSE = 0.01; “Trunk – Pelvis – LLs”: R^2^ = 0.59, RMSE = 0.01) (**Figure 6B**).

**Figure 6.**
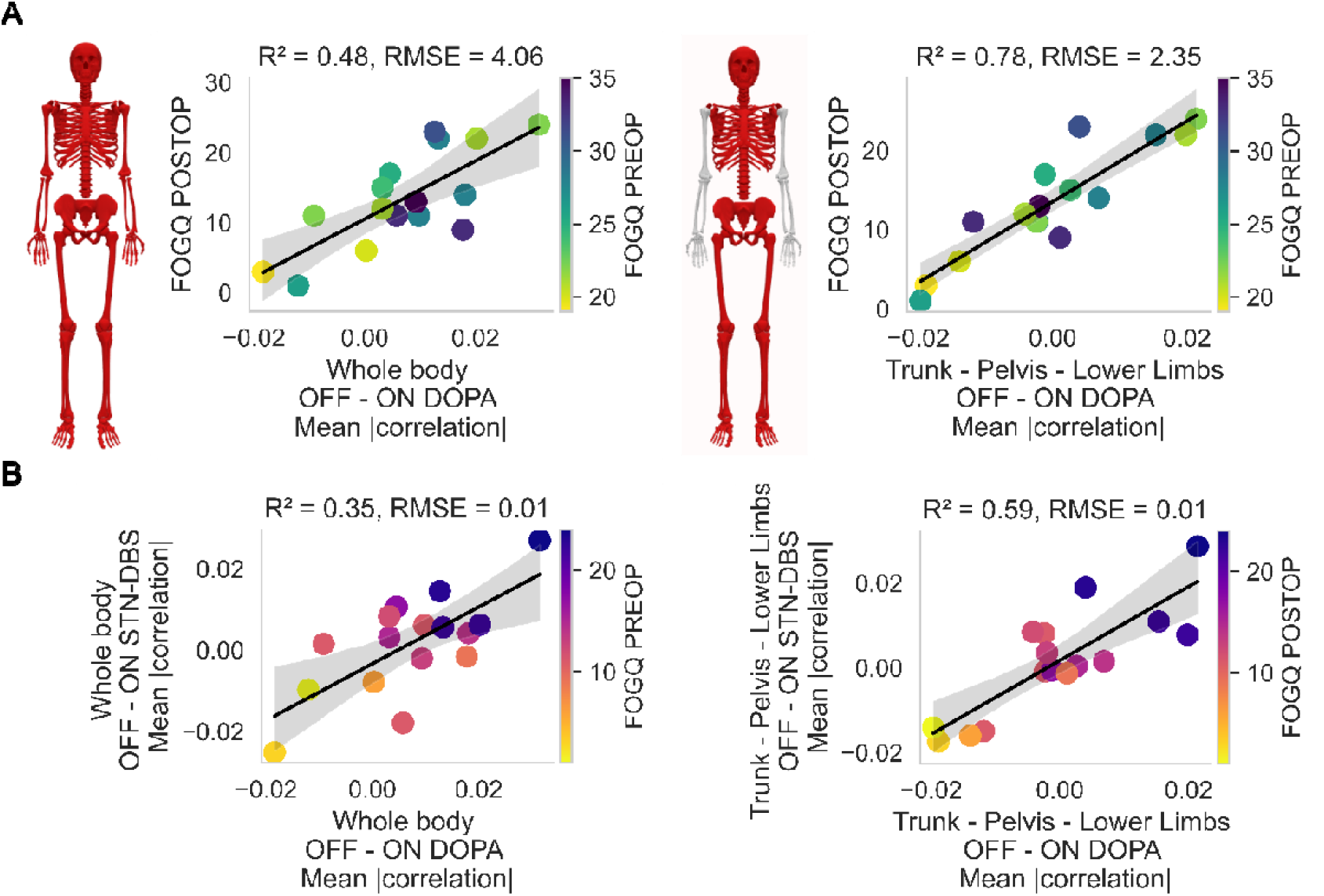
Relationship between postoperative FOG severity, the effects of STN-DBS on body-segment coordination, and preoperative DOPA effects on body-segment coordination. **A**. Scatterplot of the relationship between the POSTOP FOG-Q and PREOP DOPA-induced change in the whole-body (left) and “Trunk – Pelvis – LL” (right) coordination scores. Each dot represents one individual PD patient, color-coded by the FOG-Q PREOP (reversed *viridis* colormap). **B**. Same analysis of the relationship between the POSTOP effects of STN-DBS on these body-segment coordination scores and the PREOP effects of DOPA. Each dot represents one individual PD patient, color-coded by the FOG-Q POSTOP (reversed *plasma* colormap).

## Discussion

In this study, we assessed body-segment coordination during gait in PD patients with FOG and whether coordination metrics predict individual postoperative FOG outcomes. We found that dopaminergic medication and STN-DBS exert distinct and partially dissociable effects on gait domains and intersegmental coordination. Specific body-segment coordination patterns, rather than global spatiotemporal gait parameters, are associated with FOG severity. We showed that the preoperative coordination profile along “Trunk – Pelvis – lower limbs” axis and its modulation by dopaminergic medication predict the magnitude and direction of FOG response to STN-DBS.

Consistent with previous reports, both dopaminergic medication and STN-DBS significantly improved gait pace, reflected by increased gait speed and step length, and strengthened cortico-muscular connectivity^14,39–43^. Rhythm, however, improved only with dopaminergic medication, whereas STN-DBS did not significantly affect this domain. This dissociation supports the notion that temporal gait regulation remains partially dopamine-dependent and may rely on basal ganglia–cortical loops that are not fully restored by STN-DBS^16,44^. In contrast, dynamic balance worsened with dopaminergic medication but was preserved with STN-DBS. This finding aligns with previous observations that levodopa may worsen postural instability by increasing movement amplitude without adequately scaling postural control^18,40,45^. Conversely, STN-DBS may stabilize balance by modulating brainstem and axial control networks, possibly via downstream effects on the mesencephalic locomotor region and reticulospinal pathways^6,33^. Lastly, neither dopaminergic medication nor STN-DBS significantly altered gait variability or asymmetry. This result is consistent with prior work showing that variability-related gait features are only weakly responsive to dopamine and often persist despite optimal medical or surgical treatment, particularly in patients with FOG ^46,47^.

Although whole-body coordination remained globally unchanged across conditions, both dopaminergic medication and STN-DBS induced selective, directionally specific modifications in intersegmental coupling. In PD with FOG, reduced trunk mobility and increased arm-leg coupling have been previously reported^32,48,49^ and thought to reflect a mechanism that reduces cortical drive and stabilizes gait through interlimb entrainment when automaticity is compromised^11,15,23^. In our patients, dopaminergic medication reduced coupling between the trunk-upper limb and pelvis-upper limb while increasing coupling between the upper-lower limb and bilateral lower-limb. This pattern suggests a decoupling of upper- and lower-body coordination and a redistribution of coordination away from axial and proximal segments toward distal limb coupling under dopaminergic medication^50^. In contrast, STN-DBS predominantly increased bilateral UL coordination and certain UL-LL couplings, thereby better restoring four-limb coordination, with limited effects on trunk-pelvis interactions. This pattern aligns with previous reports of arm-swing and UL coordination normalization with STN-DBS, while axial and postural control may remain only partially improved^41,51^. Together, these findings suggest that dopaminergic medication and STN-DBS act on overlapping but non-identical components of the locomotor network, with dopaminergic medication acting mainly on the cortico-basal ganglia networks and STN-DBS acting on descending locomotor networks through direct modulation of the MLR-reticulospinal pathways^52,53^ that control spinal CPG activity and four-limb coordination patterns^54,55^.

Among all coordination features, only UL-LL coupling between the most affected UL and the least affected LL was significantly associated with parkinsonian motor disability, independent of treatment condition. This result is consistent with the link between disease severity and asymmetric interlimb coordination^12^, with the most affected UL driving this effect^25^, and impaired integration across body segments^47,48^. More interestingly, FOG severity was consistently associated with whole-body coordination, confirming the role of higher-order breakdown in postural-locomotor integration and inter-segmental coordination in FOG^19,22,31,56^.

FOG severity was substantially improved by STN-DBS. However, as previously reported, responses varied widely among individuals^38,57^. To date, reliable predictors of postoperative FOG outcomes are limited to the presence of preoperative FOG itself^33^. In this context, the identification of whole-body coordination, assessed preoperatively both OFF- and ON-DOPA, as the strongest predictor of postoperative FOG severity is novel. We also identified trunk-pelvis-LL couplings as critical predictors of postoperative FOG. These features directly reflect axial-locomotor integration, a domain previously implicated in FOG pathophysiology yet rarely quantified within a unified framework^56^. The observation that best responders exhibited coordination resembling that of healthy controls, whereas worst responders did not, supports the notion that preserved or compensable axial-lower limb coordination is a prerequisite for favorable DBS outcomes. We also observed that postoperative FOG severity and STN-DBS-induced changes in whole-body coordination correlated with dopaminergic-induced changes in this coordination, specifically in the Trunk – Pelvis – lower-limbs coordination. This suggests that coordination integrity reflects the functional state of distributed locomotor networks, including the basal ganglia, brainstem, and spinal circuits^4,6,58^, and that dopaminergic responsiveness of coordination, rather than gait speed or rhythm alone, may serve as a marker of the locomotor network’s capacity to respond to DBS. This also highlights the potential benefits of modulating spinal circuits to improve body-segment coordination and FOG^59^.

Assessing FOG severity remains a major methodological challenge in PD. FOG is episodic, context-dependent, and often inconsistently expressed during standardized assessments, and clinical scales show limited consistency^22,60^. In response to these issues, some groups aimed to develop wearable sensors to reliably quantify gait and FOG, including inertial measurement units and accelerometers placed on the trunk, pelvis, or LL^61–64^. These approaches primarily focus on event detection or local gait features rather than integrative coordination metrics and often require implementing individual algorithms to detect FOG episodes for a single patient^65^. Our findings complement these approaches by showing that FOG severity can be inferred from coordination metrics derived from a limited set of body segments, namely the trunk, pelvis, and LL. Importantly, this approach does not require the explicit observation of freezing episodes and substantially reduces the number of sensors needed. Our approach could be used for real-world daily-life assessment and monitoring of FOG severity and measures of treatment effects, addressing a critical unmet need in both clinical practice and interventional trials for FOG.

### Limitations

This study has several limitations. The sample size, particularly for postoperative analyses, was relatively small, which may limit statistical power and generalizability, although nested cross-validation and sparse modeling were used to reduce overfitting. Gait was assessed in a controlled laboratory environment, which may not fully capture the contextual variability of FOG in daily life, despite our coordination-based approach allowing inference of FOG severity even in the absence of overt freezing episodes. Finally, although informative coordination metrics were derived from a reduced set of body segments, validation using wearable sensor–based recordings is required before clinical translation.

## Conclusion

Taken together, these results highlight body-segment coordination as a clinically relevant, and potentially predictive marker of gait dysfunction and FOG in PD. Coordination metrics capture the integrative control of posture and movement, which appears central to both FOG pathophysiology and treatment response. From a clinical perspective, preoperative assessment of body-segment coordination may help identify patients at risk of poor FOG outcomes after STN-DBS and guide patient selection, counseling, and postoperative management.

## Methods

### Participants

Twenty patients with PD (14F/6M, mean age ± SD = 61.5 ± 5.1 years, body mass index (BMI) = 28.3 ± 6.8 kg/cm^2^, Table 1) scheduled for STN-DBS were recruited at Pitié-Salpêtrière and Rouen University Hospitals as part of a clinical research trial (MAGIC study, CHU ROUEN sponsor)^40^. Inclusion criteria were: 1) diagnosis of PD according to the Unified Parkinson’s Disease Society Brain Bank criteria; 2) eligibility for bilateral STN-DBS, including a > 50% improvement in motor disability with L-dopa treatment; 3) presence of severe FOG in the absence of dopaminergic medication (OFF-DOPA condition, item 2.13 of the Movement-Disorders Society Unified Parkinson’s Disease Rating Scale [MDS-UPDRS] > 0)^66^; and 4) absence of active and severe psychiatric or neurological disorders, and dementia (Mini-Mental Sate ≥ 24/30). Ten healthy controls (HCs) were also included as a reference group (GB-MOV study, INSERM sponsor, 1F/9M, mean age = 53 ± 6.5 years, BDI = 23.9 ± 2.2 kg/cm^2^)^67^.

**Table 1.** Participants demographics at inclusion.

| Patient N° | Age (y) | Weight (kg) | Height (cm) | BMI (kg/m<sup>2</sup>) | Sex | Disease duration (y) | MDS-UPDRS III OFF/ON DOPA | Axial Score OFF/ON DOPA | GABS OFF/ON DOPA | FOG-Q | MoCA | LEDD (mg/day) |
| --- | --- | --- | --- | --- | --- | --- | --- | --- | --- | --- | --- | --- |
| P01-01 | 54 | 103 | 181 | 31.4 | F | 11 | 64/14 | 20/3 | 55/9 | 22 | 26 | 1545 |
| P01-02 | 63 | 46 | 157 | 18.7 | M | 11 | 34/7 | 08/2 | 17/6 | 24 | 26 | 800 |
| P01-03 | 51 | 57 | 160 | 22.3 | F | 10 | 56/20 | 17/10 | 41/9 | 35 | 25 | 2200 |
| P01-04 | 48 | 61 | 171 | 20.9 | M | 12 | 39/9 | 12/4 | 32/7 | 33 | 30 | 1500 |
| P01-05 | 59 | 100 | 162 | 38.1 | F | 12 | 66/14 | 18/3 | 53/14 | 31 | 28 | 1300 |
| P01-06 | 65 | 64 | 169 | 22.4 | F | 11 | 68/11 | 13/2 | 39/10 | 35 | 29 | 950 |
| P02-01 | 60 | 101 | 175 | 33.1 | M | 13 | 43/10 | 11/3 | 32/4 | 25 | 26 | 1630 |
| P02-02 | 63 | 82 | 161 | 31.8 | M | 16 | 51/10 | 05/2 | 11/0 | 19 | 29 | 875 |
| P02-03 | 62 | 102 | 164 | 38.1 | M | 9 | 44/2 | 12/0 | 23/0 | 28 | 27 | 860 |
| P02-04 | 68 | 71 | 172 | 23.9 | M | 17 | 35/11 | 15/7 | 43/12 | 28 | 27 | 1725 |
| P02-05 | 58 | 91 | 171 | 31.2 | M | 6 | 34/10 | 08/3 | 26/8 | 22 | 28 | 625 |
| P02-06 | 68 | 71 | 167 | 25.6 | M | 11 | 23/06 | 06/1 | 17/1 | 21 | 28 | 2200 |
| P02-09 | 69 | 70 | 186 | 20.2 | M | 14 | 38/12 | 07/3 | 22/13 | 31 | 25 | 1464 |
| P02-10 | 63 | 112 | 181 | 34.2 | M | 14 | 47/19 | 13/3 | 31/2 | 26 | 25 | 700 |
| P02-11 | 58 | 46 | 158 | 18.4 | F | 2 | 70/21 | 10/5 | 24/9 | 28 | 30 | 500 |
| P02-12 | 56 | 78 | 182 | 23.5 | M | 10 | 45/11 | 8/0 | 16/10 | 20 | 30 | 971 |
| P02-13 | 64 | 77 | 169 | 27.0 | M | 14 | 43/14 | 09/1 | 24/4 | 33 | 25 | 2250 |
| P02-14 | 67 | 63 | 155 | 26.2 | F | 12 | 58/22 | 10/1 | 25/2 | 21 | 25 | 1286.5 |
| **Mean (SD)** | 60.9 (5.8) | 77.5 (19.6) | 168.9 (9.1) | 27.1 (6.2) | 12M/6F | 11.4 (3.4) | 47.7/12.4 (13.5/5.4) | 10.7/2.9 (4.7/2.5) | 29.5/6.7 (12.5/4.5) | 26.8 (5.2) | 27.2 (1.8) | 1258 (638) |

All studies complied with the Declaration of Helsinki and Good Clinical Practice guidelines. Ethics approvals were obtained (MAGIC trial: N° 2019-A01717-50; GB-MOV trial: N° 2012_A00225-38). Trials were registered on a clinical trial website (ClinicalTrials.gov: NCT04223427 and NCT 01682668), and all participants provided signed written informed consent.

### Gait data recordings and clinical assessment

Gait was recorded at the Paris Brain Institute Physiology and Analysis of Movement (PANAM) core facility using a three-dimensional motion capture system (200 Hz, Vicon® Oxford, UK). Thirty-two retroreflective markers were placed on anatomical landmarks following the International Society of Biomechanics (ISB) marker set (**Figure 1A**)^68^ and synchronized with a force platform (0.9x1.8 m, 1000 Hz, Advanced Medical Technology Ind, Waterform, MA, USA). Each walking trial began with the participant standing still on the force platform. Following a visual cue, participants were instructed to start walking, walk 6-8 meters at their preferred speed, make a 180° turn, and return to the starting position. An average of 20 trials were recorded per participant.

For PD patients, the severity of FOG during daily activities (with usual antiparkinsonian medication) was assessed using the FOG Questionnaire (FOG-Q)^69^, before (PREOP) and 6 months after STN-DBS (POSTOP). We also assessed the severity of gait and balance disorders and global motor disability by using the MDS-UPDRS Part III^66^ and the Gait and Balance Scale (GABS, 14 items designed to assess gait, FOG, gait cycle, balance, and posture during examination)^70^. Gait parameters, MDS-UPDRS part III and GABS scores were assessed 1) PREOP in the OFF-DOPA condition (withdrawal of 12 hours of dopaminergic treatment) and in the ON-DOPA condition (after administration of a suprathreshold dose of L-dopa), and 2) POSTOP in the OFF-DBS condition (switching OFF the DBS for one hour) and in the ON-DBS condition (usual stimulation parameters), both OFF-DOPA.

### Gait data processing

Kinematic data were preprocessed using Vicon Nexus software (version 2.14.0). Marker trajectories were filtered using a low-pass Butterworth filter with a 5 Hz cut-off frequency. Trajectory gaps shorter than 50 frames (0.25s) were reconstructed using an inter-marker correlation-based algorithm^71^.

Gait cycles were manually defined from heel strike to the subsequent ipsilateral heel strike using the vertical displacement of the calcaneus marker, and the following spatiotemporal parameters were computed using custom Matlab (R2023b, MathWorks, Natick, MA, USA) scripts: step time, swing time, stance time, cadence, step length, gait speed, and double support time.

### Quantification of body-segment coordination

To model body-segment coordination, we estimated correlations among accelerations of body markers. We measured the accelerations of 19 markers: the 7th cervical vertebra, bilateral shoulders, elbows, wrists, anterior (ASI) and posterior iliac spines (PSI), knees, ankles, heels, and toes (Figure 1A). Acceleration signals were estimated using a Savitsky-Golay filter (window length = 10 frames, polynomial order = 3). The first gait cycles were segmented, and 19x19 correlation matrices were computed to quantify inter-marker coordination along the mediolateral, anteroposterior, and vertical axes^72^ (**Figure 1B**). The mean absolute value across the three axes was computed to obtain a summary coordination matrix (**Figure 1C**). The average of the upper-triangular elements of this matrix was defined as a whole-body coordination index (**Figure 1C**). The 19 markers were then grouped into six major anatomical segments (**Figure 1D**): UL (elbow and hand), trunk (shoulders and 7th cervical vertebra), pelvis (ASI and PSI), and LL (knee, ankle, calcaneum, and toe). UL and LL were pooled according to the most affected (MA) and least affected (LA) sides based on clinical evaluation. We averaged the absolute values of pairwise correlations within each anatomical segment to yield a reduced feature space of 15 intersegmental coordination features (**Figure 1E**).

### Statistical analysis

We first assessed the effects of dopaminergic medication PREOP and STN-DBS POSTOP on gait spatiotemporal parameters and body-segment coordination. Gait parameters were aggregated by session (PREOP or POSTOP) and experimental condition (ON/OFF DOPA or ON/OFF STN-DBS). For each aggregation, mean values, variability (standard deviation), and interlimb asymmetry were computed. Asymmetry was quantified using a log-transformed ratio (*100×*|*ln(R/L)*|) averaged across right (*R*) and left (*L*) strides). After removing highly correlated parameters (*r* > 0.9, **Figure S1A-B**), 14 gait measures were retained (**Figure 2A-B**). These measures were reduced into five gait domains, *i*.*e*., *pace, rhythm, balance, variability and asymmetry*^46^, using rotated principal component analyses (RPC, **Figure 2A-B**).

For gait domain analyses across patients and trials, 143 steps were included in the OFF-DOPA condition, 237 in the ON-DOPA, 207 in the OFF-DBS, and 275 in the ON-DBS. For body-segment coordination analyses across patients and trials, 147 steps were included in the OFF-DOPA condition, 156 in the ON-DOPA, 145 in the OFF-DBS, and 155 in the ON-DBS conditions.

Comparisons of gait domains and body-segment coordination scores across conditions were performed using a paired nonparametric permutation test (10.000 permutations) for: 1) OFF-DOPA vs ON-DOPA PREOP; 2) OFF-DBS vs ON-DBS POSTOP; and 3) ON-DOPA PREOP vs ON-DBS POSTOP. False discovery rate (FDR) correction was applied using the Benjamini & Hochberg method, and statistical significance was set at p < 0.05.

Associations between FOG severity (FOG-Q) and body-segment coordination scores were evaluated using univariate Pearson’s correlation tests, both PREOP (OFF/ON DOPA) and POSTOP (OFF/ON-DBS). Given the limited sample size (n = 16) and potential non-linearities, correlations were estimated using bias-corrected and accelerated bootstrap confidence intervals with 10.000 iterations and FDR corrected. We also assessed associations between parkinsonian motor disability (UPDRS part III) and body-segment coordination scores using a linear mixed-effects model with fixed effects including the interaction between the coordination score and treatment condition (PREOP OFF/ON DOPA and POSTOP OFF/ON-DBS), and random intercepts to account for repeated measures within participants.

To identify PREOP predictors of POSTOP FOG severity, we developed a sparse linear model with LASSO regularization and a nested cross-validation framework ^73^. The model aimed to identify predictors of POSTOP FOG severity from PREOP demographics (*i*.*e*., age, disease duration, and BMI), clinical scores (i.e., FOG-Q PREOP, UPDRS III OFF/ON-DOPA, GABS OFF/ON-DOPA), gait domains (*i*.*e*., pace, rhythm, balance, variability, and asymmetry OFF/ON-DOPA), and coordination measures (*i*.*e*., whole-body coordination OFF/ON-DOPA). Model validation used an outer leave-one-out cross-validation loop to estimate generalization performance and an inner loop to tune regularization parameters. Out-of-fold predictions were aggregated, and model performance was quantified using the coefficient of determination (R^2^) and root mean squared error (RMSE) (see supplemental material, supplementary methods section). We also considered the number of times a coefficient was retained across outer folds. An additional predictive model was implemented using the same nested cross-validation framework, with body-segment coordination scores as the sole predictors of individual POSTOP FOG severity.

Following model development, we examined the relationships between 1) PREOP dopaminergic-induced changes in coordination metrics and POSTOP FOG severity, and 2) PREOP dopaminergic- and POSTOP STN-DBS-induced changes in these metrics, using leave-one-out cross-validated linear regression. Out-of-fold R^2^ and RMSE were reported.

All analyses were conducted using custom Python scripts (3.12.1; *scikit-learn, NumPy, pandas, SciPy*).

## Supporting information

Supplemental Methods and restuls

## Acknowledgments

The authors express sincere gratitude to our patients for their dedication in participating in this research. We are also grateful to Sabine Leger and Anais Hervé for their assistance in organizing the research program, and to Dorian Banier, Willy Bertucchi, Claire Olivier, and Julie Bourilhon for their help with patient inclusion and gait recordings. The authors wish to thank Fabrizio de Vico Fallani for insightful discussions during the early stages of data processing.

