## Supplemental Methods and restuls for "Body-segment coordination as a new predictor of freezing of gait in people with Parkinson’s disease"

### Supplementary material

#### Supplementary methods

The sparse linear model was inferred using a LASSO regression with the loss defined as $L(\beta) = \frac{1}{2n}||y - \beta X||_2^2 + \lambda|\beta|_1$ . We minimized the loss using scikit-learn [Pedregosa et al., JMLR 12, pp. 2825-2830, 2011]. To find the hyper-parameter in a robust fashion, we deployed a nested leave-one-out cross validation (LOOCV). This works as follows: for each patient, we create an outer validation set (composed of one patient), and an inner set of patients within which we perform LOOCV for hyper-parameter tuning. For each inner set of patients, we find the optimal $\lambda*$ through LOOCV within the inner set by minimizing the root mean squared error of the test patient across $\lambda$. Finally, using the optimal regularization parameter $\lambda*$, we infer the model on the inner set of patients and predict on the unseen validation patient. In this way, the hyper-parameter tuning is done using only data from the inner set of n-1 patients, and we obtain an outer loop prediction on the unseen validation patient. Throughout the manuscript, inference scores ($R^2$ or RMSE) were always evaluated on the outer validation patient.

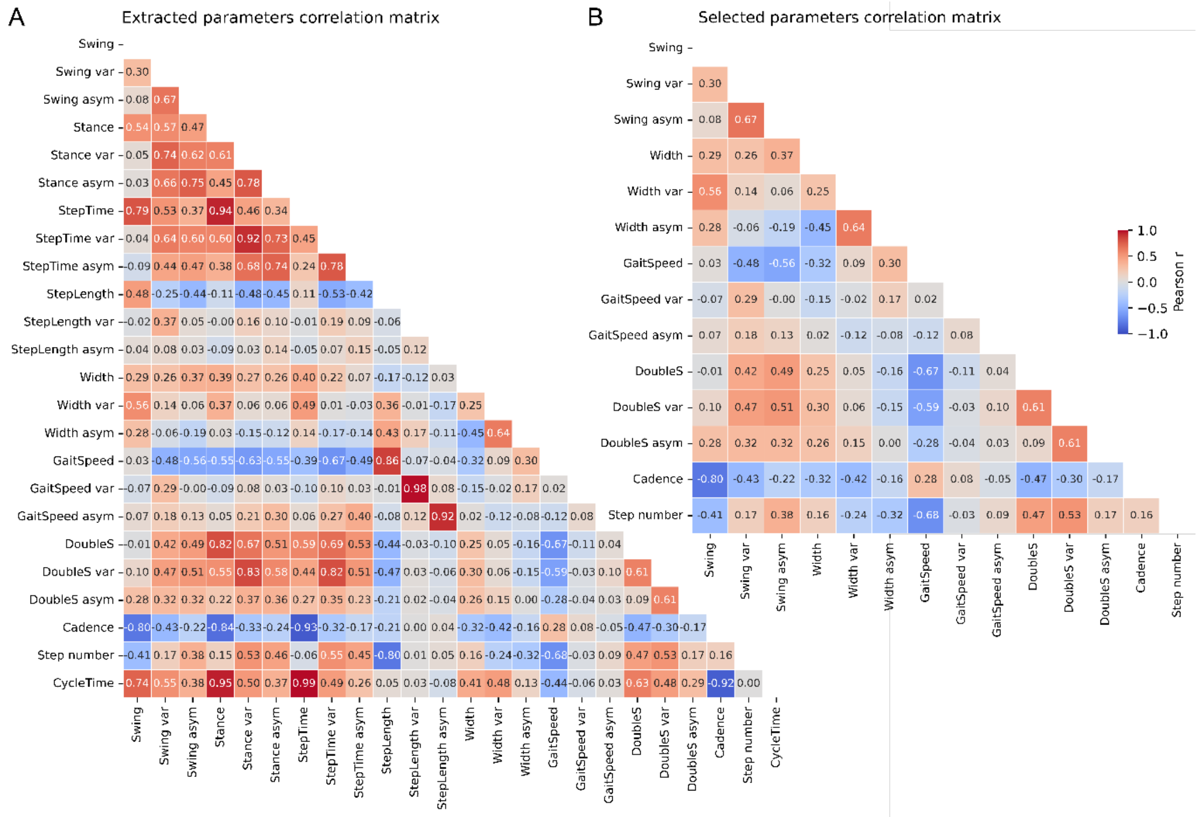


**Figure S1. Correlation matrices of spatiotemporal gait parameters.**

**A.** Correlation matrix of all extracted spatiotemporal parameters of gait cycle.

**B.** Correlation matrix for selected (non-redundant variables) used to define PCA-based gait domains. Spatiotemporal gait variables exhibiting high pairwise correlations (*r > 0.9*) were identified and excluded prior to principal component analysis (PCA). If a variable showed high correlation in any of the three derived statistics (mean, variability, or asymmetry), all three corresponding measures were removed to avoid redundancy across feature representations. This conservative approach was chosen to reduce multicollinearity and improve the interpretability and stability of the resulting principal components.

**Supplemental results**

Sixteen participants out of 20 successfully completed the study. We had two dropouts (P02-07 had to be withdrawn before randomization because of relocation abroad, which prevented the planned assessments, and P02-08 had falls with hip fracture) and two participants (P01-03 and P01-05) that could not walk independently in preoperative OFF-DOPA state.

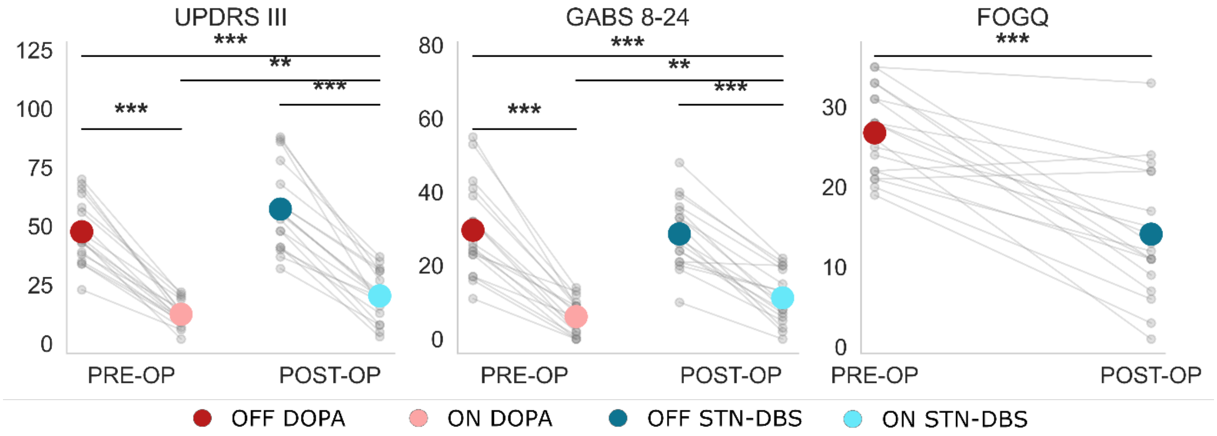


**Figure S2. Effects of DOPA and STN-DBS on parkinsonian disability, gait and balance disorders and FOG severity in PD patients.**

Graphs report the mean (big colored dots) and individual (small gray dots) scores for the UPDRS part III (left), GABS part B (middle) and FOG-Q (right) scores PREOP, OFF- (red) and ON-DOPA (pink), and POST-OP, OFF- (dark blue) and ON-STN-DBS (light blue). *p<0.05, **p<0.01, ***p<0.001 between treatment conditions.

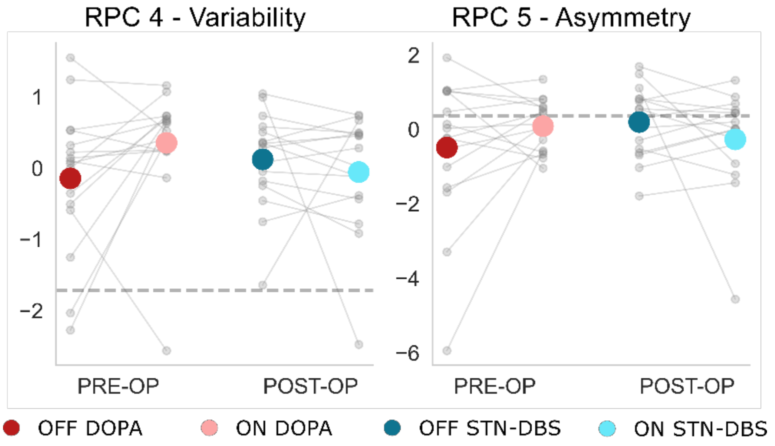


**Figure S3. Effects of DOPA and STN-DBS on variability (RPC4), and asymmetry (RPC5) scores in PD patients.**

Graphs report the mean (big colored dots) and individual (small gray dots) scores for gait *variability* (RPC4) and *asymmetry* (RPC5) PREOP, OFF-DOPA (red) and ON-DOPA (pink), and POST-OP, OFF-STN-DBS (dark blue) and ON-STN-DBS (light blue). Each dot represents one individual PD patient. The dashed grey lines represent the average of 10 healthy controls included as references. *p<0.05, **p<0.01, ***p<0.001 between treatment conditions in PD patients.

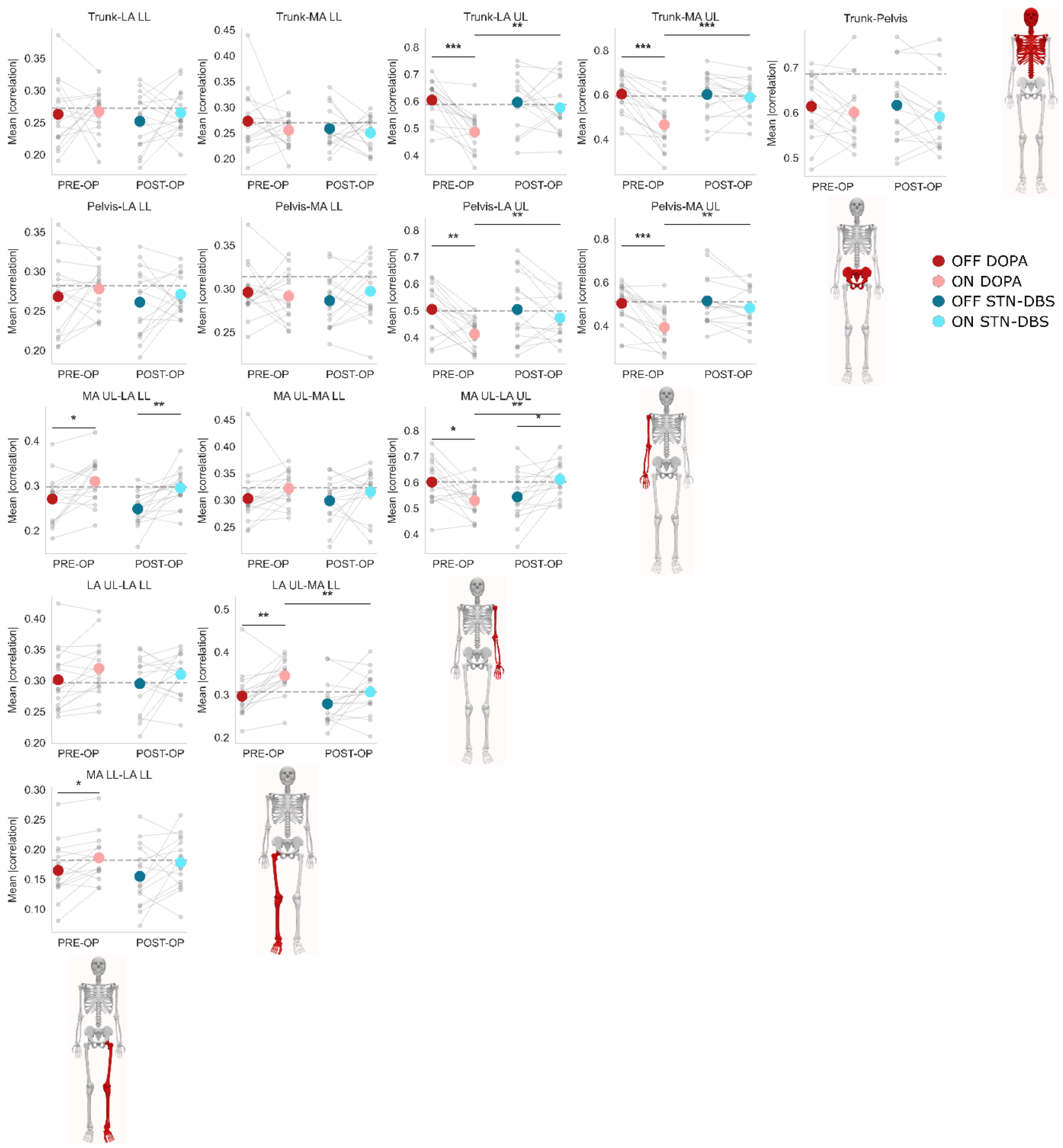


**Figure S4. Effects of dopaminergic medication and STN-DBS on body-segment coordination scores in PD patients.**

Each graph reports the mean (big colored dots) and individual (small gray dots) body-segment coordination scores PREOP OFF-DOPA (red) and ON-DOPA (pink), and POSTOP OFF-STN-DBS (dark blue) and ON-STN-DBS (light blue). The dashed gray line represents the average of 10 HC. *p<0.05, **p<0.01, ***p<0.001 between treatment conditions in PD patients.

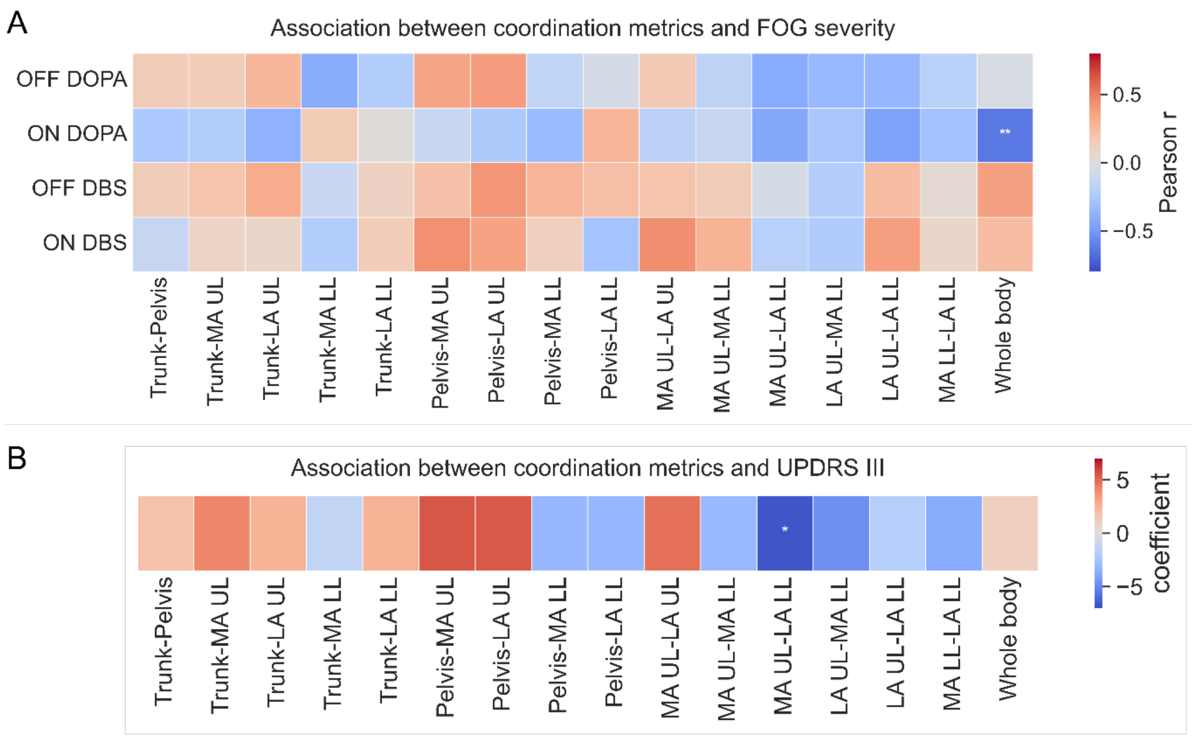


**Figure S5. Association between body-segment coordination and clinical scores.**
**A.** Heatmap of the univariate correlations between body coordination scores and FOG-Q, PREOP OFF-DOPA and ON-DOPA, and POSTOP OFF-STN-DBS and ON-STN-DBS. Bluer colors indicate negative correlations, redder colors positive ones. *p < 0.05, **p < 0.01, ***p < 0.001.
**B.** Heatmap of the univariate linear mixed effects model coefficients for body coordination scores association with MDS-UPDRS III. Bluer colors indicate negative coefficient values, redder colors positive ones. Coefficients were z-score normalized for visualization purposes. *p < 0.05, **p < 0.01, ***p < 0.001.

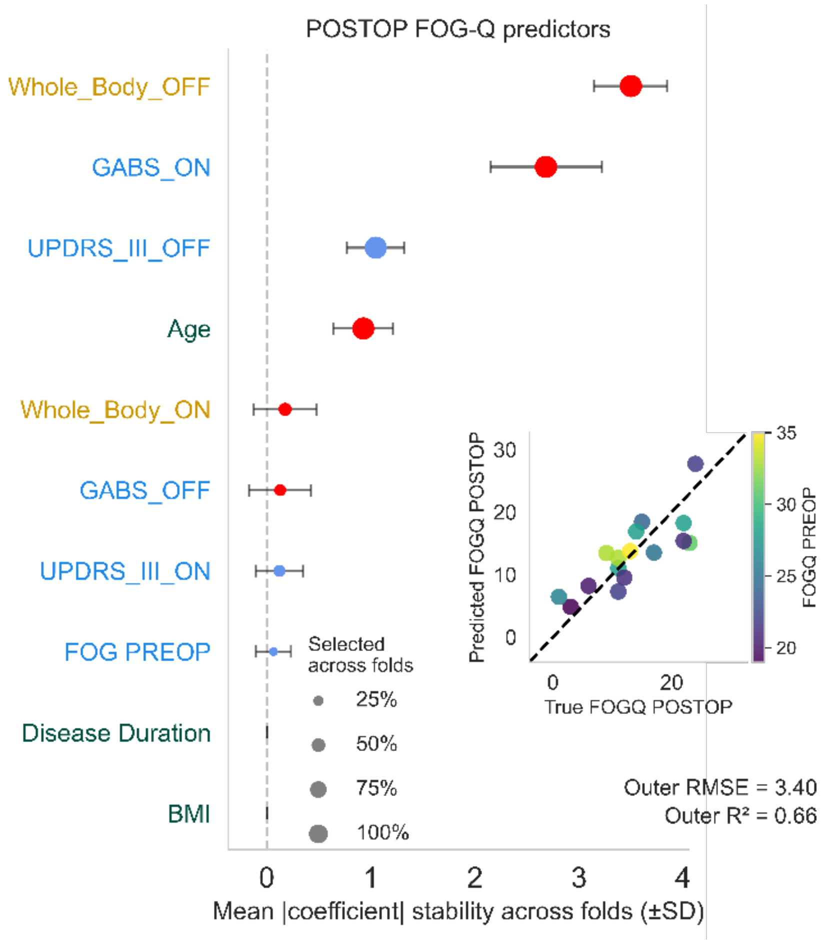


**Figure S6. Association between preoperative demographic, clinical and coordination scores and postoperative FOG severity ON-STN-DBS.**

Coefficients stability plot of the model. Dots represent the coefficients mean across each fold, and the error bars their standard deviation, and dots size the frequency of selection across folds. Red dots indicate positive coefficients and blue dots negative coefficients. Out-of-fold $R^2$ and RMSE are reported within the figure. X-axis labels are color-coded by feature type: coordination metrics (yellow), clinical scores (blue), demographics (green). The scatter plot reports the true versus predicted POSTOP FOG-Q scores ON-STN-DBS, color coded by PREOP FOG-Q scores ON-DOPA (high scores in yellow and low scores in blue); dashed black line represents identity line.
